# ENERGY EXPENDITURE DURING WALKING WITH A NOVEL TREADMILL CONTROLLER THAT INDUCES GAIT ASYMMETRY

**DOI:** 10.64898/2026.05.20.726615

**Authors:** Caitlin L. Banks, Junyao Li, Brooke L. Hall, Jan Stenum, Ryan T. Roemmich

**Affiliations:** Center for Movement Studies, Kennedy Krieger Institute, Baltimore, MD, USA; Department of Physical Medicine and Rehabilitation, Johns Hopkins University School of Medicine, Baltimore, MD, USA; Department of Neuroscience, Johns Hopkins University School of Medicine, Baltimore, MD, USA

**Keywords:** Gait analysis, walking, gait asymmetry, biomechanical phenomena, cardiopulmonary exercise test

## Abstract

Gait asymmetry is a common manifestation of walking impairment among clinical populations. We recently developed a novel treadmill walking approach called ‘dynamic treadmill walking’ that can provide asymmetric gait training by changing the treadmill speed between ‘fast’ and ‘slow’ speeds within a single stride. Here, we studied the energy expenditure associated with a variety of dynamic treadmill walking conditions. We hypothesized that the metabolic power required for dynamic treadmill walking in all conditions would approximate the metabolic power associated with conventional walking at the mean of the fast and slow speeds employed in the task. Eleven young adults without gait impairment walked on an instrumented treadmill and breathed into a metabolic measurement system. During dynamic treadmill walking, the treadmill fluctuated between 0.75m/s and 1.50m/s, each for 50% of an individual’s stride time. We used a metronome to synchronize participants’ right heel-strikes with four different timing conditions. Net metabolic power during dynamic treadmill walking was significantly greater than normal walking at the mean speed of the task (1.125m/s) and generally lower than walking at the fast speed (1.5m/s). We did not observe any significant associations between net metabolic power and several measures of gait asymmetry during dynamic treadmill walking. These findings establish dynamic treadmill walking as a promising technique for improving gait symmetry in individuals who cannot tolerate fast treadmill walking, a common gait rehabilitation approach. Future work will assess the feasibility, metabolic demands, and clinical efficacy of using dynamic treadmill walking to improve gait symmetry in clinical populations.

**New & Noteworthy:** Dynamic treadmill walking (i.e., walking with oscillating treadmill speeds) has been shown to drive gait asymmetries, but energy expenditure requirements remain unknown. We hypothesized that dynamic treadmill walking would have similar metabolic power requirements to normal walking at an intermediate speed. We found higher metabolic power requirements than the average belt speed, but less than that used for fast walking. Dynamic treadmill walking is a promising, clinically translatable technique for gait asymmetry rehabilitation.

## INTRODUCTION

Many clinical conditions can result in asymmetric gait, including stroke (1), Parkinson’s disease (2), spinal cord injury (3), orthopedic injury (4), and lower limb amputation (5). Asymmetric gait is an important target of rehabilitation because it is related to an increased risk of falling (6), an increased cost of transport (7), and decreased community engagement (8). Gait asymmetries are often heterogeneous and present in different aspects of the gait pattern – including kinematic, kinetic, and spatiotemporal parameters – depending upon the underlying neuromuscular or orthopedic deficits causing the asymmetry (1).

Several devices and therapeutic interventions exist, either in the research space or early in clinical adoption, for targeting gait asymmetries. These include specialized treadmills (9), electrical stimulation (10), robotic exoskeletons (11), and other training methods (e.g., biofeedback training and fast treadmill walking (12)). However, existing approaches have significant limitations: 1) they often require expensive, inaccessible equipment, 2) most are only capable of targeting asymmetries in specific gait parameters (e.g., step length or forward propulsion), and 3) many require significant cardiovascular fitness. There is an unmet need for new, clinically accessible approaches capable of targeting asymmetries in a variety of gait parameters, particularly those that could be implemented in patients with varying levels of fitness.

To address this need, our laboratory recently developed a ‘dynamic treadmill walking’ approach that flexibly targets asymmetries in different gait parameters by alternating the speed of the treadmill between ‘fast’ and ‘slow’ speeds during specific phases of the gait cycle as paced by a metronome (13). The metronome is used to pace heel-strikes relative to the changes in treadmill speeds because the most clinically accessible approach cannot require motion analysis or a force-sensing treadmill. Our preliminary studies in young adults without gait impairment revealed that timing of the speed changes could be customized to induce changes in symmetry of several clinically relevant gait parameters, including step length, step time, forward propulsion, and trailing limb angle (13, 14). This has demonstrated potential for development of a new, customizable intervention approach that could target different gait asymmetries using only a single-belt treadmill. However, before this approach can be tested in clinical populations (especially those with cardiovascular conditions such as stroke), it is important that we understand the metabolic demands associated with dynamic treadmill walking.

In this study, we aimed to compare the metabolic power of dynamic treadmill walking to conventional treadmill walking at constant speeds. During normal walking, metabolic power increases with increasing walking speed (15–17). However, there is some variation in the literature in other gait training approaches where the legs move at different speeds (e.g., split-belt treadmill walking). In split-belt adaptation, metabolic power first increases with larger step length asymmetries but then declines as participants adapt and gradually reduce their step length asymmetry (18). One theory for the decrease in metabolic power arises from taking advantage of the external power generated by the asymmetric treadmill that moves their legs (Sánchez *et al.*, 2019). Another split-belt walking study that mapped metabolic cost over both step length and step time asymmetries showed that change in cost is greater for step time asymmetry than step length asymmetry (19). Butterfield and Collins mapped the energy cost landscape across a variety of split-belt treadmill speeds and found that cost was generally higher than the slow belt speed and not different from the average of the two belt speeds (20). Accordingly, we hypothesized that the metabolic power associated with dynamic treadmill walking in young adults without gait impairment would approximate that required to walk at the mean of the two speeds, since 50% of each gait cycle is spent walking at each of the fast and slow speeds during dynamic treadmill walking. We considered that dynamic treadmill walking may then offer 1) customizable training for targeting asymmetries in different gait parameters, 2) an accessible approach that could be implemented by simply changing the speed of a single-belt treadmill, and 3) potential for reduced metabolic demand as compared to fast treadmill walking, a common gait training approach used in rehabilitation.

## METHODS

### Participants

A convenience sample of thirteen young adults without gait impairment (age: 24±2.9 years) were recruited from the Johns Hopkins University and the surrounding community via online announcement. Five participants self-identified as cisgender men, while eight identified as cisgender women. Five participants self-identified their race as White, six as Asian, two as Black or African American, and all identified as not Hispanic or Latino. All participants reported to be right leg dominant. Technical difficulties required exclusion of two participants’ data (one for improper mask seal after a rest break and one for metabolic cart software malfunction). The data presented in this manuscript therefore include eleven participants (age: 24±3 years, 5 male/6 female, 5 Asian/2 Black/4 White).

Inclusion criteria included individuals that had no major orthopedic or neurologic impairments, could walk on a treadmill for at least ten consecutive minutes, and could tolerate walking while wearing the metabolic testing equipment. Exclusion criteria included advanced age, inability to walk on a treadmill for ten consecutive minutes, or major medical conditions that impair walking or make study procedures challenging. No potential participants were excluded from participating in the study. The Johns Hopkins School of Medicine Institutional Review Board 2 approved all study procedures (IRB00143977). Participants provided written informed consent, and the study conformed to the standards set by the Declaration of Helsinki, except for registration in a database. The authors attest that our study procedures and reporting are in compliance with the *Journal of Physiology* human ethics policy. Participants were provided a small monetary compensation for their time.

### Study protocol

Participants walked at three Baseline speeds for five minutes each: 0.75m/s, 1.125m/s, and 1.50m/s (Figure 1). Next, they walked with the dynamic treadmill controller engaged for six ten-minute trials. The first and last trials with the controller were open-loop and unconstrained (i.e., we instructed participants simply to walk with the treadmill as it fluctuated between 0.75m/s and 1.50m/s to determine whether they would select a preferred pattern). Participants then completed four *Metronome* trials (*Fast, Slow, Accelerate, Decelerate*) in a randomized order. During these trials, we instructed the participants to time their right heel-strike with the tone of a metronome, which was timed according to that individual’s average stride time as measured during the first minute of the 1.125m/s Baseline trial. The metronome allows control of heel-strike timing to the four different conditions listed above without requiring additional instrumentation (e.g., motion capture or treadmill force plates) that would not be accessible in a clinical setting. This is a replication of the Experiment 2 protocol from our previous study (13), with the addition of metabolic testing.

**Figure 1.**
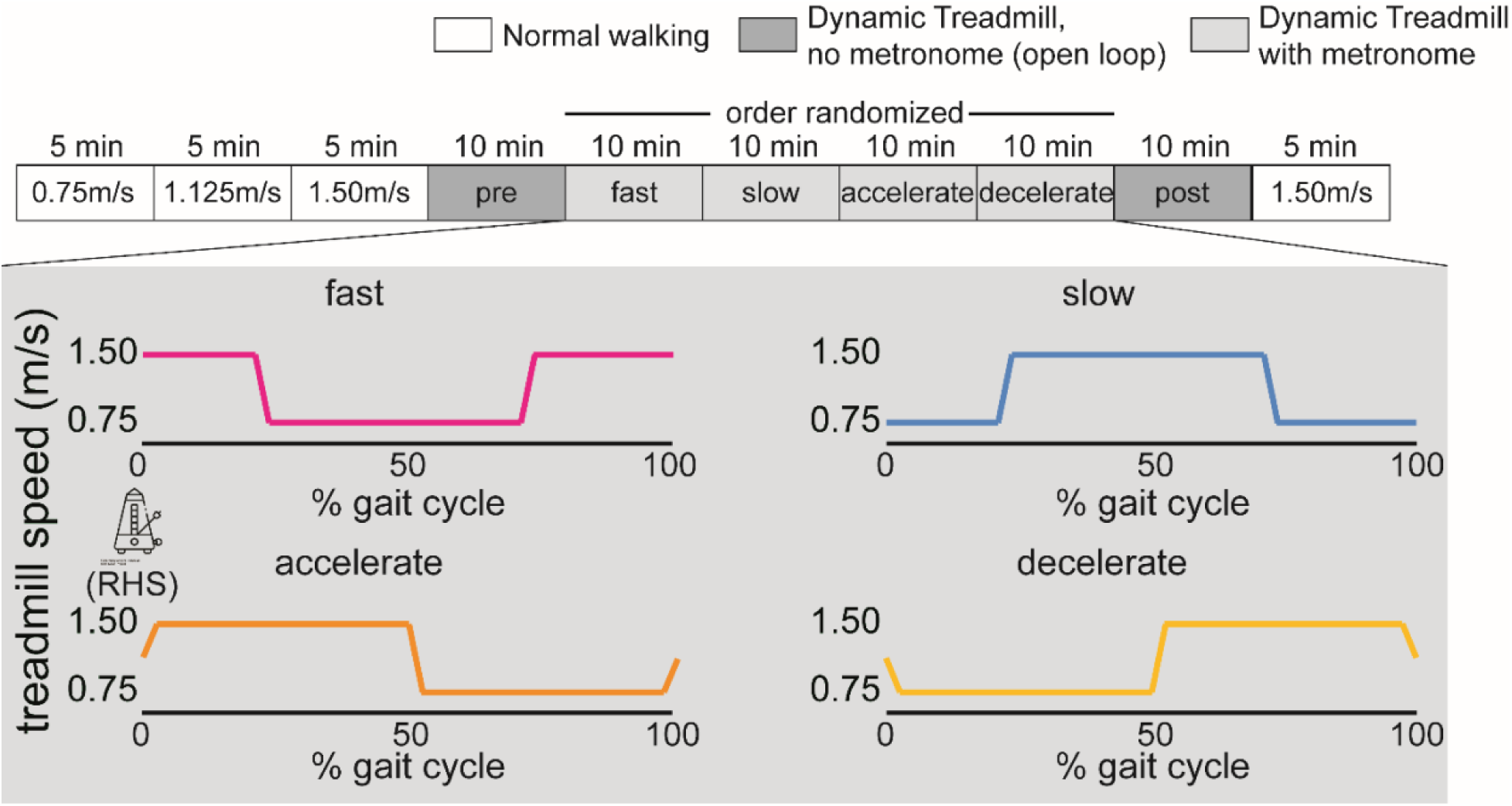
Experimental protocol. A timeline of each treadmill walking condition is depicted along the top row of the figure. Initially, participants walked normally at three fixed speeds for five minutes each: 0.75m/s, 1.125m/s, and 1.50m/s. The next six trials, depicted in grey boxes, represent the dynamic treadmill walking trials. Of those six dynamic treadmill trials, there were two open-loop trials (dark boxes, labelled ‘pre’ and ‘post’) where there was no metronome pacing the right heel-strikes. The four light grey shaded trials had right heel-strikes paced to a metronome, with each condition’s treadmill speed depicted below the timeline. Conditions are named for the speed of the treadmill at right heel-strike (i.e., at fast speed, at slow speed, accelerating to fast speed, decelerating to slow speed). Figure modified from Browne et al. Fig 1B <u>(2023)</u>. Metronome image by Andi Nur Abdillah for Noun Project (CC BY 3.0).

The dynamic treadmill controller runs the treadmill via a custom script utilizing the D-Flow (Motek Medical, Amsterdam, NL) programming environment. The treadmill fluctuates between 0.75m/s and 1.50m/s with an acceleration rate of 6.0m/s^2^. Within each gait cycle, the controller changes speed at time intervals corresponding to 50% of each individual’s stride time from the Baseline 1.125m/s trial. In the *Metronome* trials, the software plays an audible beep to the pace the participant’s right heel-strike as follows:

- *Fast* condition: right heel-strike paced to occur at the fast treadmill speed, resulting in the fast treadmill speed (1.50m/s) for 25% of the gait cycle, slow (0.75m/s) for the middle 50%, and fast for the remaining 25%.
- *Slow* condition: right heel-strike paced to occur at slow treadmill speed, inverse of *fast* condition.
- *Accelerate* condition: right heel-strike paced to occur while the treadmill at approximately 1.125m/s, when treadmill was accelerating between the slow and fast speeds. This results in the first 50% of the gait cycle at the fast speed (1.125m/s) and the second 50% at the slow speed (0.75m/s).
- *Decelerate* condition: right heel-strike paced to treadmill decelerating from fast (1.125m/s) to slow (0.75m/s) speed, inverse of *accelerate* condition.

Following the six dynamic treadmill controller trials, participants completed another five-minute trial of conventional walking at 1.50m/s for comparison of metabolic data before and after dynamic treadmill walking.

### Data collection

Participants walked on an instrumented split-belt treadmill (Motek Medical, Amsterdam, NL) surrounded by a nine-camera three-dimensional motion capture system (Vero, Vicon Motion Systems, Centennial, CO, USA; 100 Hz; RRID: SCR_015001). Participants always wore a fall-arrest harness that did not provide body weight support when walking on the treadmill. The treadmill has emergency stop buttons located near each study team member, and light gates at the front and back of the treadmill that stop the belts so participants cannot walk off either end. Passive retroreflective markers were placed according to the Vicon lower body Plug-in Gait marker set with the addition of bilateral fifth metatarsal heads, medial malleoli, medial femoral epicondyles, greater trochanters, iliac crest, and trunk markers placed approximately at C7, T10, jugular notch, and xiphoid process (21–23). Participants wore their own walking shoes and fitted athletic clothing. Three-dimensional ground reaction forces were collected at 1000 Hz using the treadmill’s built-in force plates under each belt.

Metabolic data were collected using a TrueOne 2400 system (Parvomedics, Sandy, UT, USA). We sampled breath-by-breath oxygen consumption and carbon dioxide production. Baseline metabolic rate was measured during two minutes of quiet standing.

### COVID-19 protocol

Most of the data collection for this study was conducted in the fall and winter of 2022-2023. Because expired air from the metabolic chamber was released into the room air, we took additional precautions to lower the risk of exposure to COVID-19 for our participants and study staff. We ran a high efficiency particulate air purifier (Coway Airmega, Seoul, South Korea) on the highest setting throughout the duration of the experiment, placed next to the expired air outlet. Study staff self-screened for respiratory illness symptoms and did not come into the building if they were feeling ill. Staff were fit tested for N-95 masks by the Johns Hopkins Medicine Respiratory Protection Program and wore them throughout the data collection. Participants took a Flowflex COVID-19 rapid antigen test (ACON Laboratories, San Diego, CA, USA) after being consented and prior to removing their facemask and being set up for metabolic testing. Finally, ‘Do Not Enter – Metabolic Experiment in Progress’, signs were placed on lab doors to minimize traffic and contact with non-study personnel. No participants declined to follow our procedures, and no participants tested positive on the day of study enrollment.

### Data analysis

We used custom MATLAB (The Mathworks, Natick, MA, USA; RRID: SCR_001622) software for data analysis. We filtered marker data using a fourth order low-pass Butterworth filter with a cut-off frequency of 6 Hz. We determined heel-strike and toe-off events as the timings of the maximum and minimum values of the limb angle trajectory (defined as the sagittal angle between a vector drawn from the iliac crest marker to the second metatarsal head marker and a vector from the iliac crest to the ground) (24). We calculated spatiotemporal and kinematic variables for comparison to our previous work and correlation with metabolic power. Information on how these metrics are calculated can be found in our previous publication (13). Average values from the last 30 strides of each trial are included in all statistical analyses.

Due to difficulties with treadmill software recording belt speeds, we had to calculate belt speed using marker data for six participants. To confirm accuracy of our calculation method, we calculated speeds for the remaining trials and compared to the recorded belt speeds, yielding peak speed root mean square errors of 0.03±0.04m/s.

We calculated metabolic power using Brockway’s equation derived for energy expenditure (25). We then calculated net metabolic power by subtracting baseline metabolic power (from a static standing trial) and normalizing to each participant’s body mass. Finally, we calculated summary values over the last two minutes of each gait trial once participants had reached a metabolic steady state. Technical difficulties with the treadmill computer required excluding the final minute of *Decelerate* trial for one participant: accordingly, we calculated metabolic power over minutes 7-9 for this participant.

### Statistical analysis

We conducted all statistical analyses in R version 4.3.1 (R Core Team, 2023; RRID: SCR_001905) via RStudio version 2023.06.1 (Posit Team, 2024; RRID: SCR_001905). We utilized the following R packages for all analyses: tidyverse (RRID: SCR_019186), readxl (RRID: SCR_018083), ggplot2 (RRID:SCR_014601), plyr (RRID:SCR_026985), svglite, desctools (RRID:SCR_027454), and dplyr (RRID:SCR_016708) (28–34). We assessed net metabolic power data for normality using the Shapiro-Wilk W test and data were normally distributed (*W*=0.98, *p*=0.19). We also tested net powers for homogeneity of variance using Bartlett’s test (K^2^=4.53, df=9, p=0.87). We utilized repeated measures ANOVA with a within-subjects factor of Trial to compare metabolic power across the different walking conditions using the rstatix package (35). We then conducted post-hoc pairwise comparisons using two Dunnett’s tests, where we compared net metabolic power during the dynamic treadmill conditions against the metabolic power measured by normal walking at the average of the two dynamic treadmill belt speeds (1.125m/s) and at the fast belt speed (1.50m/s). For validation purposes, we also performed a repeated measures ANOVA on the three Baseline speeds alone, compared pre-post differences in the 1.50m/s conditions using a two-tailed t-test, and repeated our repeated measures ANOVA using block order rather than grouping by dynamic treadmill condition to assess potential order effects. We then assessed the association between net metabolic power and spatiotemporal, kinematic, and kinetic measures of gait symmetry using linear regression. Because the open loop condition did not show consistent changes in gait symmetry in the initial study, we maintained it in the study protocol but do not focus our results on the open loop conditions. We set α≤0.05 for all analyses.

## RESULTS

### Speed profiles of dynamic treadmill walking

Figure 2 shows the average speed profiles in each dynamic treadmill metronome condition. To verify that participants stepped in synchrony with the metronome in the paced dynamic treadmill trials, we computed cross correlation coefficients between the actual treadmill speed profile and the expected speed profile for each condition, anchored to each right heel-strike. A one-way repeated measures ANOVA revealed no significant differences in cross-correlation values for the last 30 strides of each metronome trial (F(3,24)=1.65, *p*=0.21, η^2^=0.11). While there are no statistical differences, the speed profiles show that participants tended to be better at performing the Fast and Slow conditions than the Accelerate and Decelerate conditions.

**Figure 2.**
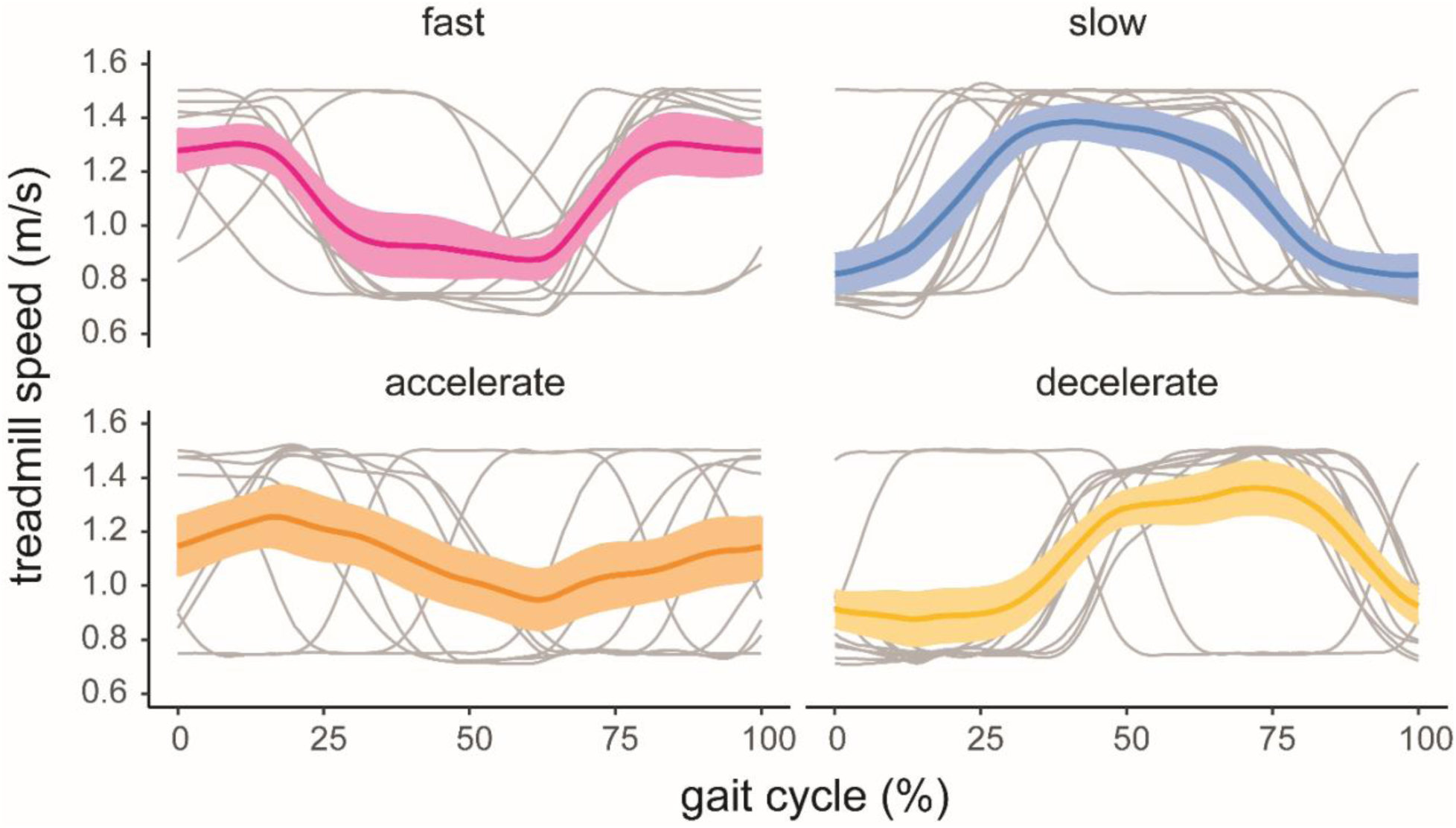
Treadmill speed profiles for each metronome trial. Each participant line (gray) represents the average treadmill speed profile over the last 30 strides of each trial, while the colored lines and shading represent group mean and standard error.

### Metabolic power

Net metabolic power for each participant, as well as averaged across all participants, is shown in Figure 3. We conducted a repeated measures ANOVA to assess the effects of Trial on net metabolic power. We observed a significant main effect of Trial on net metabolic power (F(9,90)=52.42, *p*<0.001, η^2^=0.66). Pairwise comparisons using Dunnett’s test compared the six dynamic treadmill trials to the average belt speed (Baseline 1.125m/s) as the control trial. All dynamic treadmill trials showed significantly higher net metabolic power than the Baseline 1.125m/s trial (all *p*’s<0.01). We completed a second Dunnett’s test to compare the dynamic treadmill trials to the 1.50m/s Baseline trial, which resulted in the Baseline 1.50m/s trial requiring higher metabolic power than the Accelerate (p=0.04) and Decelerate (p=0.03) trials and not being different from the remaining four trials (p’s>0.05). Table 1 shows each of the statistical comparisons and the associated 95% confidence intervals.

For methods validation purposes, we also performed a repeated measures ANOVA on the three Baseline speeds, tested potential fatigue effects by comparing pre-post differences in the 1.50m/s conditions, and repeated our analysis above using trial completion order rather than grouping by dynamic treadmill condition to assess potential order effects. Net metabolic power in the Baseline trials scaled with walking speed (there was a main effect of Trial: F(2,20)=190.14, *p*<0.001, η^2^=0.77), with all pairwise comparisons reaching statistical significance (*p’s*<0.001). Net metabolic power in the Post 1.50m/s condition was greater than in the Baseline 1.50m/s condition (t(10)=2.77, *p*=0.02, mean difference=0.35W/kg), suggesting presence of a fatigue effect following more than 60 minutes of treadmill walking. Finally, we did not observe a significant order effect, as there was no systematic pattern of change in net metabolic power across the metronome conditions in the order that they were tested (Supplemental Figure 1). A repeated measures ANOVA showed that trial number was significantly related to net metabolic power (F(9,90)=53.08, *p*<0.001, η^2^=0.66). Pairwise comparisons revealed no differences between all pairings of dynamic treadmill trials (all p’s>0.05).

**Figure 3.**
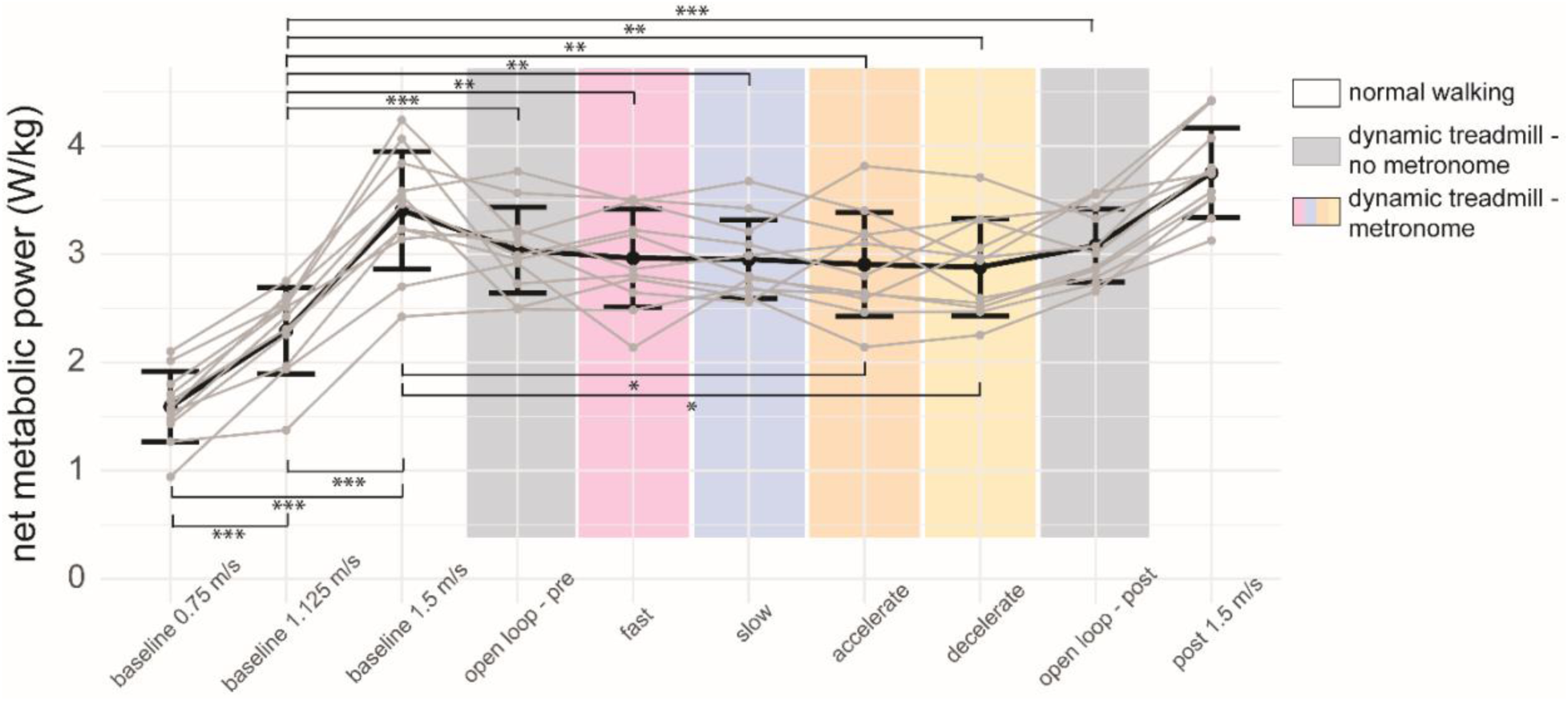
Net metabolic power differs across walking trials. Net metabolic power during dynamic treadmill walking is higher than the average belt speed but lower than fast treadmill walking. The dynamic treadmill trials with gray shading represent open loop conditions without a metronome, while the colored shading represents the metronome-paced trials. Thin grey lines represent individual subject data, while black lines represent group average ± standard deviation. Statistically significant pairwise comparisons are noted by asterisks: *: p<0.05; **: p<0.01; ***: p<0.001.

**Table 1.** **Multiple comparisons via Dunnett’s test reveal differences between all dynamic treadmill trials and Baseline walking at the average belt speed** (1.125m/s, shaded region), as well as differences between select dynamic treadmill trials and Baseline walking at the fast belt speed (1.50m/s, unshaded).

| Comparison | Mean Difference | Confidence Interval | p-value |
| --- | --- | --- | --- |
| **Open Loop-Pre vs. Baseline 1.125m/s** | **0.746** | **(0.282, 1.210)** | **0.000** |
| **Fast vs. Baseline 1.125m/s** | **0.671** | **(0.207, 1.135)** | **0.002** |
| **Slow vs. Baseline 1.125m/s** | **0.660** | **(0.197, 1.124)** | **0.002** |
| **Accelerate vs. Baseline 1.125m/s** | **0.613** | **(0.149, 1.076)** | **0.005** |
| **Decelerate vs. Baseline 1.125m/s** | **0.588** | **(0.124, 1.052)** | **0.007** |
| **Open Loop-Post vs. Baseline 1.125m/s** | **0.786** | **(0.323, 1.250)** | **0.000** |
| Open Loop-Pre vs. Baseline 1.50m/s | -0.367 | (-0.857, 0.123) | 0.22 |
| Fast vs. Baseline 1.50m/s | -0.442 | (-0.932, 0.048) | 0.09 |
| Slow vs. Baseline 1.50m/s | -0.452 | (-0.942, 0.038) | 0.08 |
| **Accelerate vs. Baseline 1.50m/s** | **-0.500** | **(-0.990, -0.010)** | **0.04** |
| **Decelerate vs. Baseline 1.50m/s** | **-0.523** | **(-1.015, -0.035)** | **0.03** |
| Open Loop-Post vs. Baseline 1.50m/s | -0.326 | (-0.816, 0.164) | 0.31 |

### Biomechanics of dynamic treadmill walking

Figure 4 illustrates a selection of gait parameters that induced asymmetries (step length, trailing limb angle, and peak propulsion) in select *Metronome* trials. We chose this subset because they are relevant to common neurologic gait asymmetries, although all variables shown in Browne et al. were consistently reproduced (13).

**Figure 4.**
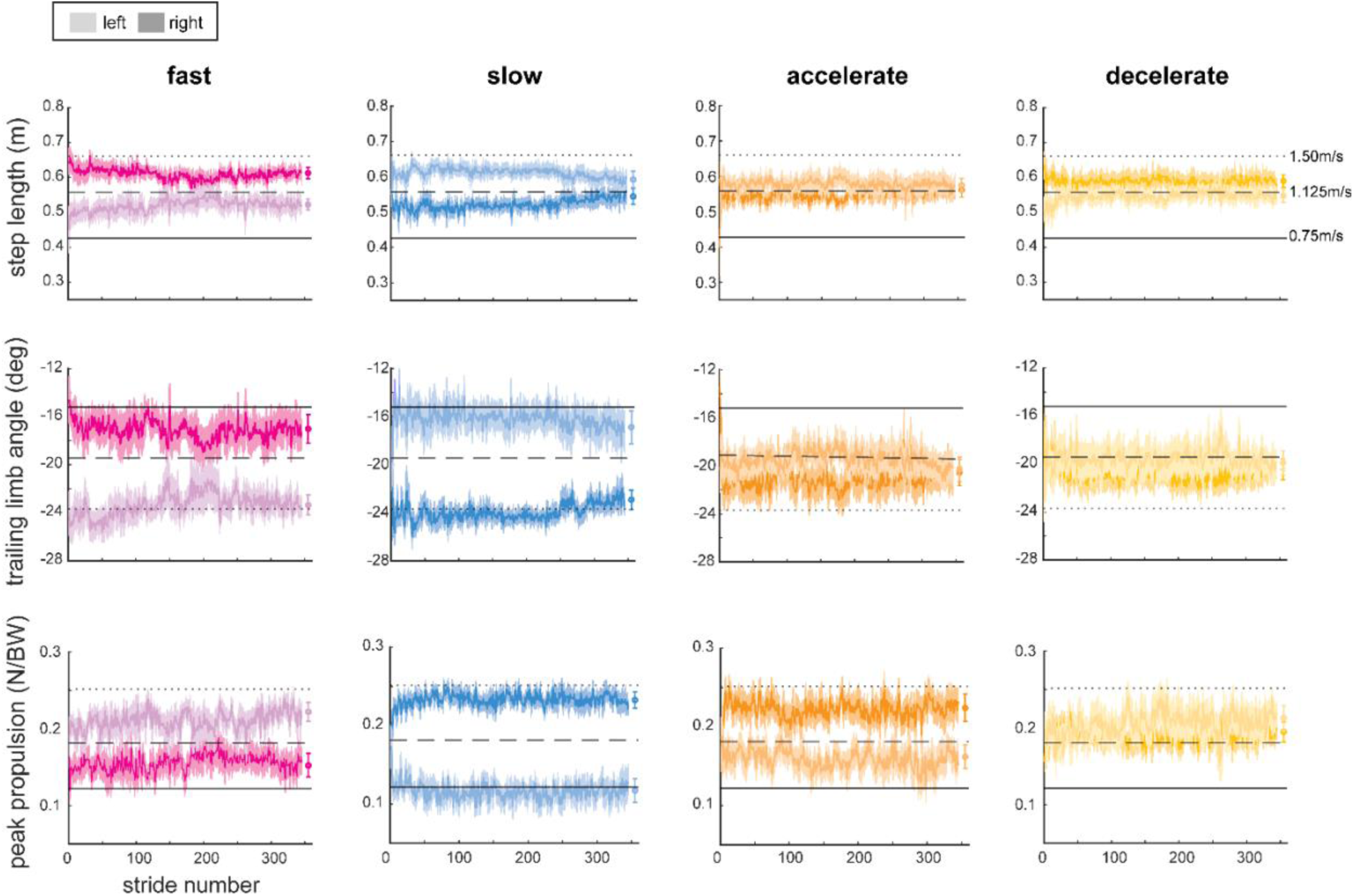
Spatiotemporal, kinematic, and kinetic asymmetries agree with our previously published work. Here, we show select biomechanical variables from the four metronome trials, as labelled in the top row, but the gait parameters across all trials agreed with prior work (13). Step-by-step values of step length (top), trailing limb angle (middle), and peak propulsion (bottom) are illustrated in each plot (group mean ± standard error of the mean), with overall mean values from the last 30 strides depicted on the end of each time series. Solid horizontal lines indicate normative values for the Baseline 0.75m/s trial, dashed for Baseline 1.125m/s, and dotted for Baseline 1.50m/s.

### Relationship between metabolic power and biomechanics

We were interested in understanding what factors explain the added cost of dynamic treadmill walking, and whether selected biomechanical asymmetries induced by dynamic treadmill walking were associated with net metabolic power within each *Metronome* condition. We performed linear regression models with net metabolic power as the dependent variable and three select asymmetry values of interest from our prior work as potential predictors: step length asymmetry, trailing limb angle asymmetry, and peak propulsion asymmetry (Figure 5). None of these parameters – step length asymmetry (*F*(4,39)=0.08, *p*=0.99, *R^2^_adj_*=-0.09), trailing limb angle asymmetry (*F*(4,39)=0.23, *p*=0.92, *R^2^_adj_ =*-0.077), or peak propulsion asymmetry (*F*(4,39)=0.085, *p*=0.99, *R^2^_adj_ =*-0.093) – were significant predictors of net metabolic power.

## DISCUSSION

Dynamic treadmill walking creates a novel, asymmetric walking pattern with metabolic power demand that falls between normal walking at the two belt speeds employed in the task. This walking pattern requires more metabolic power than the mean belt speed for normal walking, which aligns with expectation that walking with gait asymmetries is more costly than a typical, symmetric walking pattern, and generally less metabolic power than the fast walking speed. The timing of belt changes relative to stepping (i.e., the metronome condition) does not significantly change the metabolic power required to complete the task. Increasing net metabolic power with increasing baseline walking speed agrees with the literature and our understanding of human physiology (15, 16). As a comparison against another asymmetric gait training approach, dynamic treadmill walking appears to require slightly more metabolic power than split-belt treadmill walking. The metabolic cost of split-belt walking is greater than slow belt normal walking, lower than fast belt normal waking, and not different from the average of the two belt speeds once a steady-state pattern is reached (18, 20).

**Figure 5.**
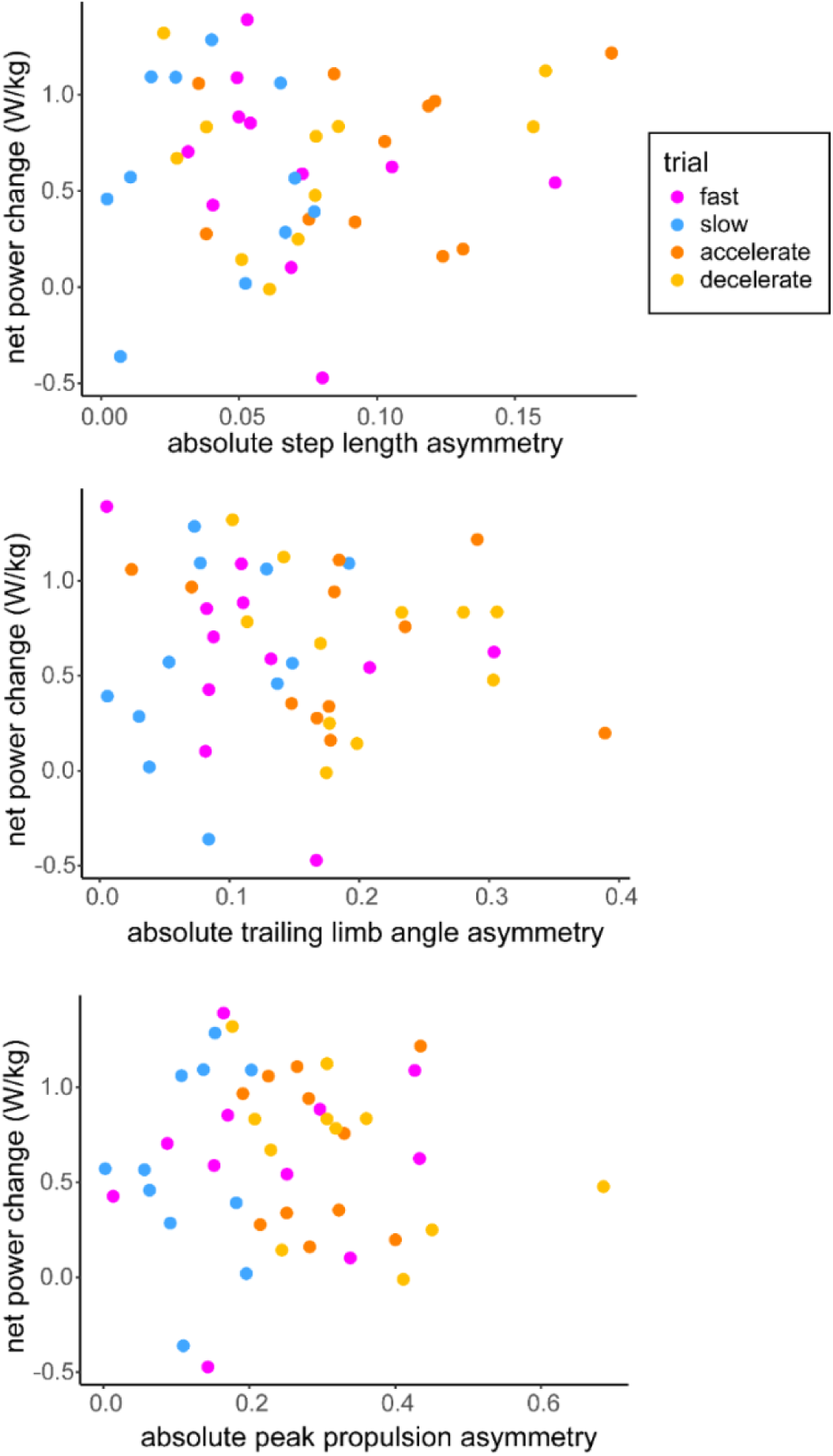
Change in net metabolic power is not associated with biomechanical asymmetries in this sample. Scatterplots show net metabolic power change (difference between trial net power and Baseline 1.125m/s net power; y-axis) vs. the absolute value of: step length asymmetry (top), trailing limb angle asymmetry (middle), and peak propulsion asymmetry (bottom) averages for each participant from the last 30 strides of the trial. Linear regression analysis showed that none of these asymmetry parameters were significant predictors of net metabolic power change.

Dynamic treadmill walking appears to present a more explicit response to stepping to the metronome and having the legs moved by the treadmill belts than split-belt treadmill walking. This can be seen in Figure 4 with relatively steady biomechanical parameters over several minutes of walking and in our previous work by the lack of robust aftereffects during washout trials (13). It is worth noting that we did not introduce the belt speed changes gradually in this paradigm. Instead, we started each of the dynamic treadmill trials with the belts immediately fluctuating between the high and low speed limits. It is possible that a more gradual introduction to belt speed changes, coupled with the closed-loop controller tested in our previous study (13), could induce more of an expected motor adaptation pattern. However, a closed loop controller is less clinically feasible than the open-loop, metronome paced conditions. Although we used a split-belt treadmill with custom control software to develop the protocol and measure kinematic and kinetic patterns associated with the task, the technology itself could be feasibly adapted for clinical use. A former team member created a dynamic treadmill using a store-bought treadmill and an off-the-shelf controller board, and future technology development will continue after demonstration of the clinical feasibility within populations with gait asymmetry. While we require the laboratory technology to perform thorough biomechanical characterization of the behavioral response, the goal of this controller has always been translational feasibility.

Our observation that dynamic treadmill walking requires less metabolic power than normal walking at the faster belt speed, as well as our finding that none of the dynamic treadmill conditions differed from each other, is particularly helpful from a rehabilitation perspective. One of the standard evidence-based physical therapy recommendations for improving post-stroke gait is fast walking (36). While fast walking can improve walking speed and has some evidence for improving gait biomechanics (12), it is not specifically targeted to improving gait symmetry. However, it is not necessarily safe or tolerable for some individuals with neurological conditions to walk continuously at fast enough speeds to improve gait biomechanics. Conversely, techniques like split-belt treadmill walking (37), circular treadmill walking (38), and robotic orthoses (39) can modify gait symmetry, but require specialized equipment that is not available in most rehabilitation clinics. Like split-belt walking, individuals walking on the dynamic treadmill only need to experience the fast speed for a portion of each gait cycle. However, this dynamic treadmill approach can offer asymmetric gait training without the need for a split-belt treadmill. Our dynamic treadmill controller offers the opportunity for targeted, participant-specific asymmetric gait modification without the metabolic demand of fast walking. This could be particularly useful for individuals that are deconditioned after a hospital stay or have underlying cardiac conditions that make fast walking too taxing.

There have been few studies investigating dynamic treadmill walking, so there are some limitations to be addressed in future work. This study included a small sample of young, unimpaired adults with a limited range of both net metabolic power and biomechanical variables of interest across participants, which may relate to why we did not find correlations between metabolic power change and biomechanical asymmetries. Further characterization would be necessary to understand the mechanism behind the increased metabolic power requirements and their impact on the resulting biomechanical asymmetries. Additionally, we do not yet know how individuals with gait impairment or individuals substantially older or younger than this group would respond to the task. Due to the novelty of the task, our dynamic treadmill findings have not yet been replicated independently. Finally, we do not yet know what multiple exposures to the controller will do to an individual’s gait. Studies are ongoing to assess how individuals with gait impairment interact with the dynamic treadmill. We recently characterized how dynamic treadmill walking changes with different belt speeds and how the induced asymmetries alter interlimb biomechanics (14).

Our next steps are to 1) assess the metabolic demands and biomechanical patterns of dynamic treadmill walking in participants with gait asymmetry post-stroke, and 2) assess the efficacy of training with participant-specific asymmetry reduction. We are also conducting additional studies in young adults without gait impairment to gain a better understanding of the mechanisms driving asymmetries created by dynamic treadmill walking.

## CONCLUSIONS

This study represents the first assessment of the metabolic demands of dynamic treadmill walking. We found that the asymmetric walking pattern created by the dynamic treadmill controller requires more metabolic demand than normal walking at the average treadmill belt speed. This metabolic power requirement is also higher than what is reported in the literature for split-belt walking. While dynamic treadmill walking is more energetically costly than the average belt speed, it does require less metabolic power than normal walking at the faster belt speed. This has potential for use as a clinically translatable training mechanism for individuals with gait asymmetry that cannot tolerate fast walking.

## ACKNOWLEDGEMENTS

This research was supported by the American Heart Association (grants #935556 and 26BCDA1622699, RTR) and the National Institute of Child Health and Human Development (P50HD118624, RTR, JS). CLB was supported by the Kennedy Krieger Institute and Johns Hopkins University School of Medicine Research Training in Rehabilitation for Brian Injury and Neurological Disability program (NICHD grant #2T32HD007414-31, PI Bastian), followed by an American Heart Association Postdoctoral Fellowship (https://doi.org/10.58275/AHA.25POST1366391.pc.gr.227425). BLH was supported by a Research Supplement to Promote Diversity in Health-Related Research (#3R21HD110686-02S1, PI RTR) and is currently supported by a Ruth L. Kirschstein Institutional Predoctoral Training Program in the Neurosciences (5T32NS091018-26, PI Knierim). BLH and CLB implemented data science and visualization training in this work from the ReproRehab program (NIH NICHD/NCMRR R25HD105583, PI Liew).

## ADDITIONAL INFORMATION

### Data Availability Statement

All data are individually illustrated within the figures and tables. Data files and analysis code, as well as supplementary figures, are archived on Zenodo: https://doi.org/10.5281/zenodo.20293357.

### Author Contributions

**Conceived and designed research:** CLB, JS, RTR

**Performed experiments:** CLB, BLH, JL

**Analyzed data:** CLB, BLH

**Interpreted results of experiments:** CLB, BLH, JS, RTR

**Prepared figures:** CLB, BLH

**Drafted manuscript:** CLB, JL, BLH

**Edited and revised manuscript:** CLB, JL, BLH, JS, RTR

**Approved final version of the manuscript:** CLB, JL, BLH, JS, RTR

### Author Pronouns

**She/her/hers:** CLB, BLH

**He/him/his:** JL, JS, RTR

### Current Addresses

**Junyao Li:** Feinberg School of Medicine, Northwestern University, Chicago, IL, United States

### Authors Eligible for Early Investigator Prize

CLB, BLH

