## Supplemental Figure 1 for "ENERGY EXPENDITURE DURING WALKING WITH A NOVEL TREADMILL CONTROLLER THAT INDUCES GAIT ASYMMETRY"

**SUPPLEMENTAL MATERIAL**

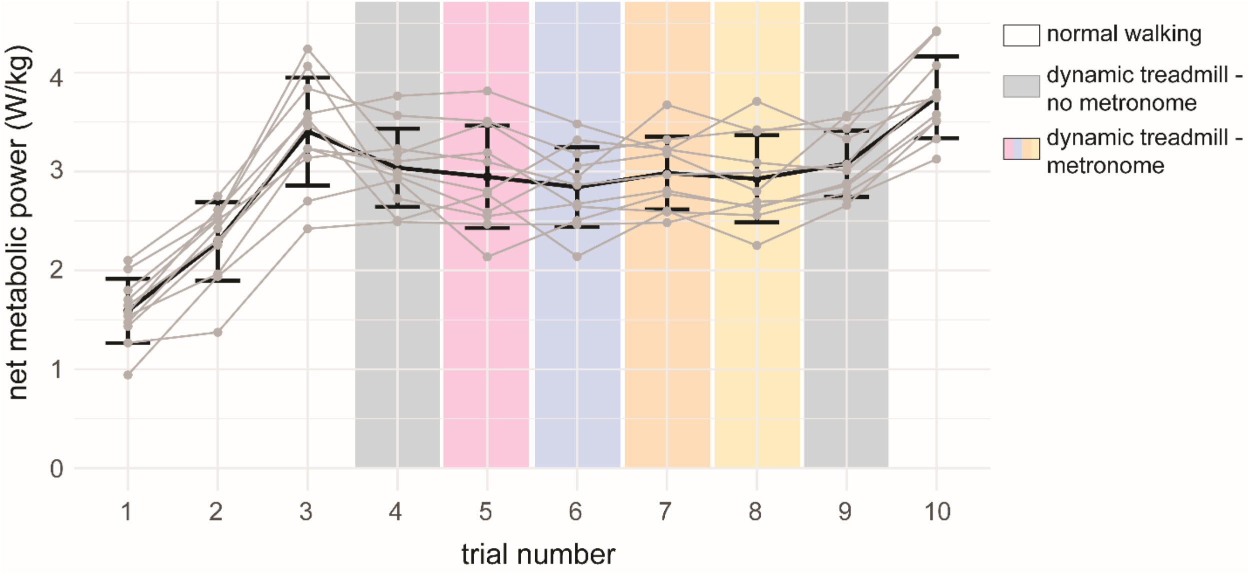


**Supplemental Figure 1.** When arranged by testing order, there are no order effects for net power during metronome trials from those in Figure 3. Testing order of the four metronome-paced trials (Blocks 5-8) was randomized within each participant, order of all other Blocks is consistent with protocol from Figure 1. A repeated measures ANOVA showed that Block was significantly related to net metabolic power ($F(9,90)=53.08$, $p<0.001$, $\eta^2=0.66$). Thin grey lines represent individual subject data, while black lines represent group average ±1 standard deviation.
